# *Mycobacterium* phage Butters-encoded proteins contribute to host defense against viral attack

**DOI:** 10.1101/2020.04.27.053744

**Authors:** Catherine M. Mageeney, Hamidu T. Mohammed, Marta Dies, Samira Anbari, Netta Cudkevich, Yanyan Chen, Javier Buceta, Vassie C. Ware

**Affiliations:** Department of Biological Sciences, Lehigh University, Bethlehem, PA 18015; Department of Chemical and Biomolecular Engineering, Lehigh University, Bethlehem, PA 18015; Department of Bioengineering, Lehigh University, Bethlehem, PA 18015

## Abstract

A diverse set of prophage-mediated mechanisms protecting bacterial hosts from infection has been recently uncovered within Cluster N mycobacteriophages. In that context, we unveil a novel defense mechanism in Cluster N prophage Butters. By using bioinformatics analyses, phage plating efficiency experiments, microscopy, and immunoprecipitation assays, we show that Butters genes located in the central region of the genome play a key role in the defense against heterotypic viral attack. Our study suggests that a two component system articulated by interactions between protein products of genes *30* and *31* confers defense against heterotypic phage infection by PurpleHaze or Alma, but is insufficient to confer defense against attack by the heterotypic phage Island3. Therefore, based on heterotypic phage plating efficiencies on the Butters lysogen, additional prophage genes required for defense are implicated.

**IMPORTANCE:** Many sequenced bacterial genomes including pathogenic bacteria contain prophages. Some prophages encode defense systems that protect their bacterial host against heterotypic viral attack. Understanding the mechanisms undergirding these defense systems will be critical to development of phage therapy that circumvents these defenses. Additionally, such knowledge will help engineer phage-resistant bacteria of industrial importance.

## INTRODUCTION

Mycobacteriophages – viruses infecting mycobacterial hosts-are of interest because they are useful in diagnostics of mycobacterial infections (1), the most notable of which is tuberculosis (TB), and additionally can serve as genetic tools for mycobacteria (2-5). Most recently, engineered mycobacteriophages have been used in therapeutic applications to combat infections from antibiotic-resistant strains of *Mycobacterium abscessus* (6). To date over 11,000 mycobacteriophages have been isolated, over 1,800 sequenced, and over 1,600 are available in GenBank (7, 8). Mycobacteriophages are a small subset of the estimated 10^31^ bacteriophages existing in the biosphere (9). Mycobacteriophages display high levels of genetic diversity and have been divided into 29 genomically similar clusters (A-AC) and a group of singletons with no close relatives (10, 7). Although an increase in isolation and genomic characterization of mycobacteriophages has occurred recently, the void in knowledge about gene expression and function of mycobacteriophage gene products remains.

Prophages make up a majority of the known bacteriophage population (11). The relationship between prophages and bacterial strains has shown numerous benefits to both the hosts and phages. Prophages confer many advantages to the host upon integration such as enhanced fitness, reduction of mutation rates, selective advantages, and defense against additional viral attack (12). The bacterial host in turn provides the phages with protection from harsh environments (12). In this context, numerous mechanisms of defense have been recently discovered for *Pseudomonas, Mycobacterium, and Gordonia* prophages (13-16), with the expectation that prophage-mediated defense systems are likely widespread throughout the bacterial-phage world. Cluster N phages have been investigated for defense mechanisms that allow the host bacterium to become resistant to heterotypic phage attack (14). Currently 29 Cluster N mycobacteriophage genomes are found in GenBank (8). Cluster N mycobacteriophages are characterized by small genomes (40.5-44.8kbp) (14; phagesdb.org). These bacteriophages have siphoviridae morphologies and are capable of integration into the *Mycobacterium smegmatis* mc^2^155 *attB* site tRNA-Lys (MSMEG_5758) (17, 14).

Here we focus on *Mycobacterium* phage Butters that was isolated on *M. smegmatis* mc^2^155. Butters is one of the smallest members of Cluster N with a genome of 41,491bp (18) and contains 66 open reading frames (ORFs). The Butters genome can be divided into three regions (Figure S1). Genes in the first region are rightward-transcribed, encoding structural genes such as capsid and tail proteins (genes *1-25*). The central portion of the genome (genes *26-40*) encodes two endolysins (Lysin A and Lysin B), a holin, genes used for integration and excision of the genome, and, importantly, many genes with unknown functions. Within the central region of all Cluster N genomes is the “variable region” (Figure S1) that has considerable genomic variation among all Cluster N phages (14). Finally, the third region includes rightward-transcribed genes (genes *41-66*) encoding proteins used in DNA maintenance and many of unknown function.

Cluster N mycobacteriophage prophage-mediated defense is a function of genes in the central variable region (14). Genes *30* and *31*, are in the Butters variable region and were originally classified as orphams (i.e., genes with no known mycobacteriophage counterpart) prior to their discovery in a recently characterized Cluster N phage Rubeelu. Yet, their function remains unknown. These genes are among those expressed in a Butters lysogen (14), rendering them as suitable candidates that mediate defense of the lysogen against heterotypic phages.

Here we used bioinformatic analyses, heterotypic phage plating efficiency experiments, microscopy, and immunoprecipitation experiments to explore the roles of gp30 and gp31 in protecting a Butters lysogen from phage attack. Our results suggest that gp30 and gp31 interact and that gp31 may have an impact on the subcellular localization of gp30. Efficiency of plating data on *M. smegmatis* strains expressing gp30, gp31, or gp30 and gp31 combined, show that PurpleHaze attack is completely abolished when gp30 is expressed alone but infection is partially restored when gp30 is co-expressed with gp31. Moreover, for Cluster A9 mycobacteriophage Alma, viral attack is significantly inhibited by gp30, but no inhibition is observed when gp30 is co-expressed with gp31. Altogether, we propose that gp30-gp31 interaction is instrumental against specific viral attack. Further, since the proposed Butters gp30/gp31 system has no apparent effect on attack by Cluster I1 phage Island3, we suggest a gp30-independent defense mechanism against this phage. Collectively, these data demonstrate that multiple defense mechanisms are encoded by the Butters prophage.

## RESULTS

### Bioinformatics analyses predict transmembrane domains (TMDs) for Mycobacteriophage Butters gp31 but not for gp30

Butters gp30 (GenBank protein ID: AGI12977.1) and gp31 (GenBank protein ID: AGI12978.1) were analyzed for transmembrane domains using TMHMM (19, 20). Butters gp30 was not predicted to have any TMDs (Figure 1A), while gp31 is predicted to have four (Figure 1B). Two additional proteins, gp28 and gp21 (GenBank protein IDs: AGI12968.1; GI12975.1, respectively), were analyzed by TMHMM and used as bioinformatics controls. A known membrane protein, gp28 (annotated holin) is predicted to have two TMDs (Figure S2A) and an annotated minor tail protein, gp21, has no predicted hydrophobic domains, suggesting its cytoplasmic localization (Figure S2B). These results are indicative of cytoplasmic localization for gp30 and membrane integration for gp31.

**Figure 1.**
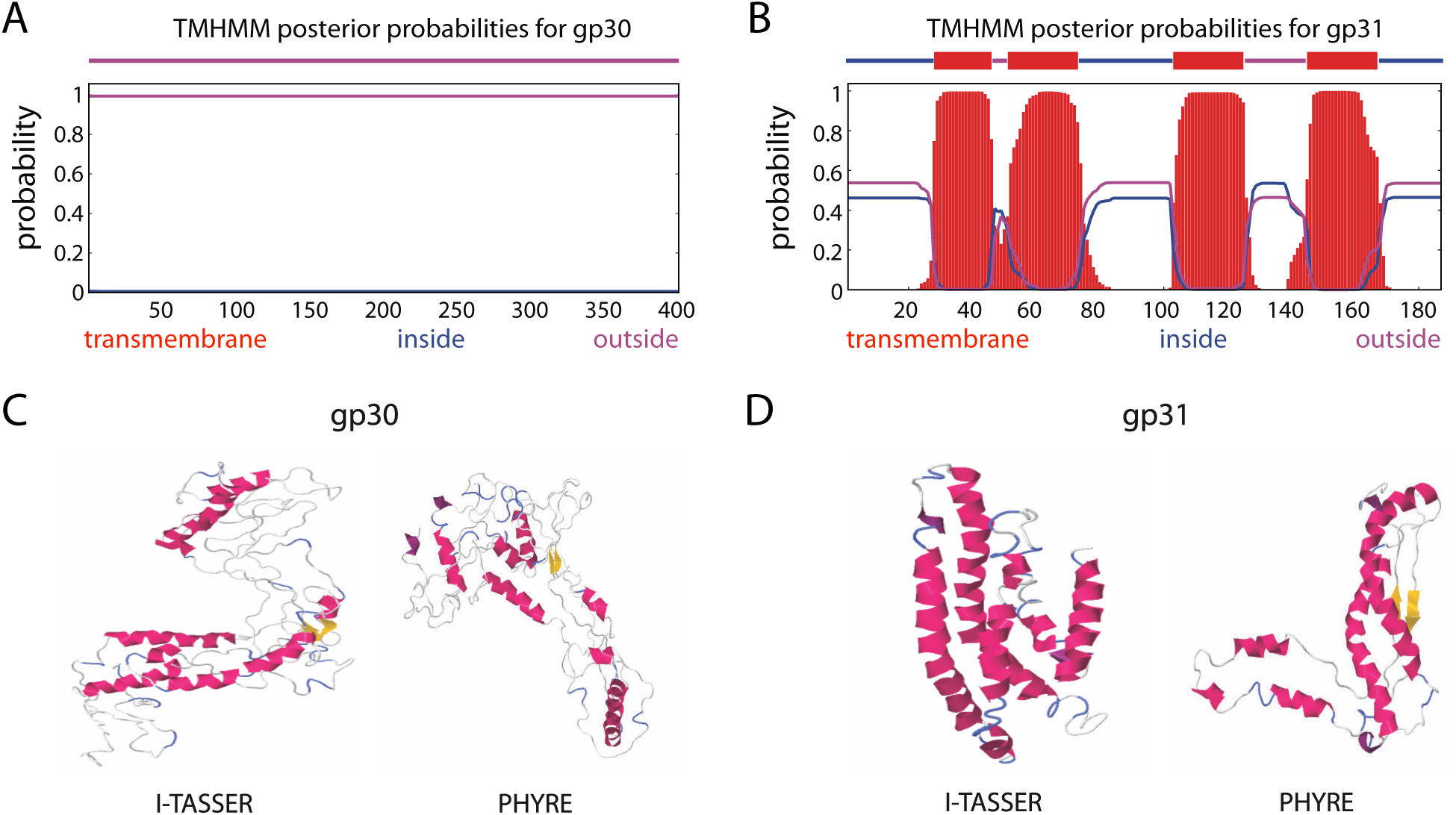
Posterior probabilities for protein gp30 (**A**) and gp31 (**B**) as predicted by TMHMM (19, 20). The amino acids index is shown on the horizontal axis. The blue, purple, and red lines indicate the probability of an aa being located inside, outside, or within the cell membrane, respectively. Butters gp30 is predicted as a protein with domains outside the membrane (cytoplasmic). Butters gp31 is predicted to have 4-pass transmembrane domains (membrane protein). **C, D.** Predicted secondary structures of proteins gp30 (**C**) and gp31 (**D**) using I-TASSER (21) and PHYRE (22). The long, parallel, alpha helices of gp31 are characteristic of membrane proteins as predicted by TMHMM.

I-TASSER (21) and PHYRE (22) were used to further analyze gp30 and gp31 structures. Gp30 has weak homology with protein structures in the PDB and no distinguishing features (Figure 1C). Butters gp31 is predicted to have 4 alpha-helices which presumptively are membrane spanning in concord with the TMHMM posterior probabilities for gp31 (Figure 1D).

Gp30 and gp31 were also analyzed using HHpred to investigate function (23, 24). HHpred analysis of gp30 yields a weak hit to the motif DUF4747 (Probability: 69.48, E-value: 140) (Figure 2A). This DUF4747 domain is conserved in the cytoplasmic components of the Abi systems uncovered in coliphage Lambda [RexA] (25, 26), *Mycobacterium* phage Sbash [gp30] (15), and *Gordonia* phage CarolAnn [gp44] (16) (Figure 2B). Lambda cytoplasmic RexA (when activated by a protein-DNA complex of the invading phage) binds to the membrane protein RexB (an ion channel) which depolarizes the membrane resulting in loss of intracellular ATP, death of the bacterium, and abortion of infection (27). Similar mechanisms of action have been proposed for the Abi systems of Sbash (15) and CarolAnn (16). Remarkably, Butters gp31 and all the membrane components of these Abi systems have 4 transmembrane domains (Figure 1 and Figure S3). These findings highlight the possibility that Butters gp30 and gp31 may play roles in prophage-mediated defense in a way analogous to the RexAB Abi system. Butters gp31 has weak homology to bacteriophage holins from Enterobacter phage P21 (probability: 58.8, E-value: 25), *Haemophilus* phage HP1 (probability: 52.88, E-value: 39), pneumococcal phage Dp-1 (probability: 21.24, E-value: 550), and to a bacteriophage holin family, superfamily II-like (probability: 64.23, E-value: 26) (28). However, it is atypical for holin proteins to have more than two TMDs (29). Moreover, gene *31* is expressed in the Butters lysogenic cycle (14), rendering a holin function unlikely for gp31.

**Figure 2.**
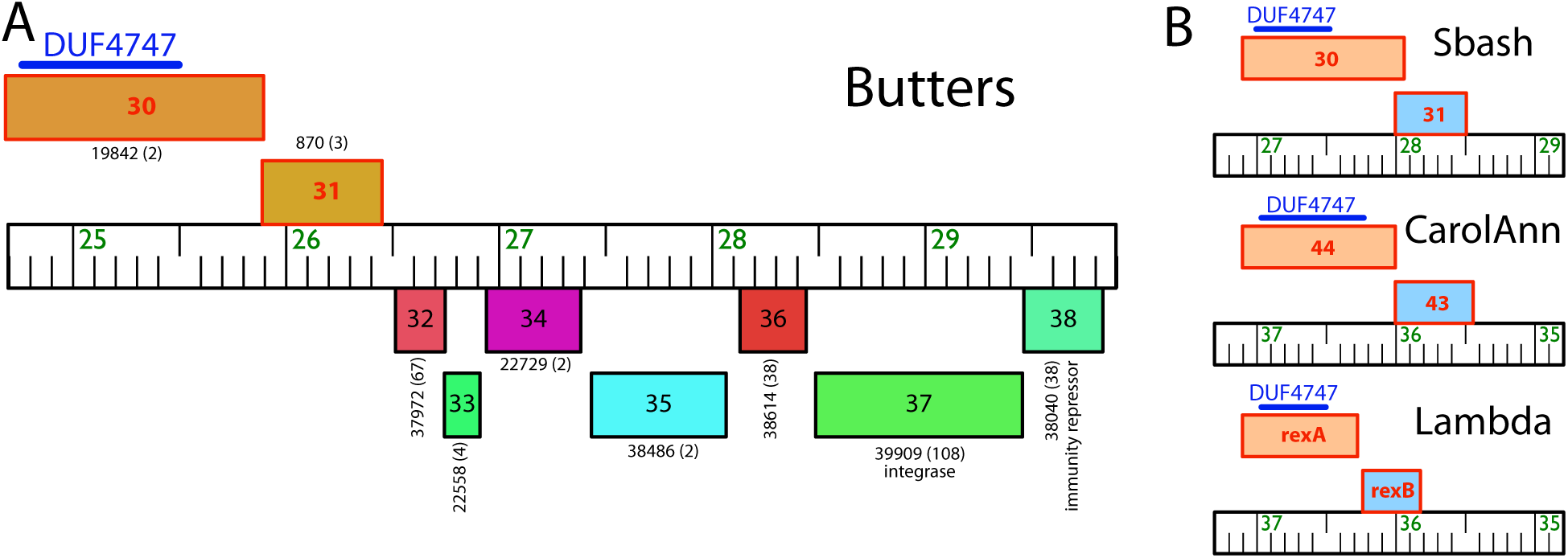
Genomic synteny of selected phage-encoded exclusive systems **A.** Central “variable region” of Butters genome. The gene colors and numbers represent gene phamilies designated by Phamerator database Actino_Draft (version 353) (40); the number of phamily members is shown in parentheses. Rightward and leftward transcribed genes are shown above and below, respectively. The blue bar on top of gene *30* indicates the DUF4747 domain. **B.** Syntenic representation of two-component exclusion systems found in bacteriophages Sbash, CarolAnn, and Lambda. Butters genes *30* and *31* are compared to the Abi systems of Sbash, CarolAnn, and Lambda. Genes (represented as boxes) are aligned to their genome (ruler) labeled with coordinates. The conserved DUF4747 domain is aligned on the putative cytoplasmic component of the exclusion system (blue bar). Transcription is from left to right in all cases. Genomes of CarolAnn and Lambda have been reversed to aid comparison.

### Phage infection assays indicate that gp30 and gp31 are components of a prophage-mediated defense system against viral attack

Given the shared structural homology between Butters gp30 and gp31 and the Abi systems of coliphage lambda, *Gordonia* phage CarolAnn, and Mycobacteriophage Sbash (Figures 2 and S3) coupled with the fact that all characterized Cluster N mycobacteriophage prophage-mediated defenses have been mapped to genes within the central variable region of their genomes (14), we hypothesized that Butters genes *30* and *31* are involved in prophage-mediated defense. We tested this hypothesis using a phage infection assay. We spotted serial dilutions of a selected panel of heterotypic phages known to be inhibited by the Butters lysogen: Alma and Island3 (14; this study) and PurpleHaze (this study) on lawns of *M. smegmatis* mc^2^155 derivatives expressing Butters gene *30* alone, Butters gene *31* alone, and both Butters genes *30* and *31* represented as mc^2^155(gp30), mc^2^155(gp31), and mc^2^155(gp30-31), respectively (Figure 3). Phage serial dilutions were also spotted on a Butters lysogen, mc^2^155(Butters), and a Butters lysogen variant with gene *30* deleted, mc^2^155(ButtersΔ*30*).

**Figure 3.**
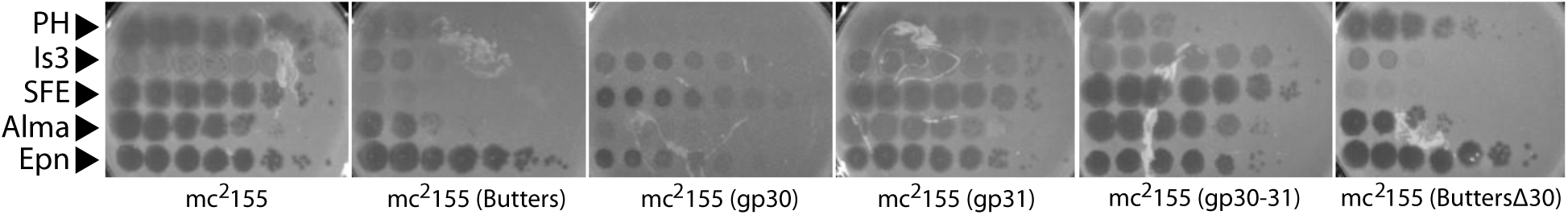
Plating efficiencies of heterotypic phages on *M. smegmatis* mc^2^155 strains expressing gp30, gp31, or gp30-31 (designated as mc^2^155(gp30), mc^2^155(gp31), and mc^2^155(gp30-31), respectively). Phages spotted are listed on the left: PH (PurpleHaze); Is3 (Island3); SFE (ShrimpFriedEgg); Alma; Epn (Eponine). Phage lysates were serially diluted to 10^−7^ and spotted (3µL each) onto a lawn of each bacterium plated with 1xTA. ShrimpFriedEgg (Cluster N) inhibition on mc^2^155(Butters), and mc^2^155(ButtersΔ*30*) is repressor-mediated (14). mc^2^155(gp30) defends against PurpleHaze(A3) and Alma(A9) but not Island3(I1). gp30-mediated defense is attenuated in the presence of gp31. In agreement with previous results (14), Island3 and Alma show reduced plating efficiencies on mc^2^155(Butters). On both lysogen lawns, the absence of individual plaques in the dilution series for Island3 and ShrimpFriedEgg suggest that observed clearings are due to “killing from without” and not infection.

All phages efficiently infected an *M. smegmatis* mc^2^155 strain carrying the empty vector pMH94 (Figure S4A). Eponine(K4) plated efficiently on all lawns while ShrimpFriedEgg(N) was inhibited by the lysogenic strains expressing the Butters immunity repressor (Figure 3 and Table S1). Heterotypic phages PurpleHaze(A3), Island3(I1), and Alma(A9) had reduced efficiency of plating on an *M. smegmatis* mc^2^155(Butters) lawn (14; Figure 3 and Table S1). Defense against heterotypic phages is independent of immunity repressor function (14); therefore, we would predict that inhibition of PurpleHaze, Island3, and Alma infection would be mediated by other genes. *M. smegmatis* mc^2^155 strains expressing Butters gp30 alone, completely abolished PurpleHaze infection, reduced infection of Alma by 4 orders of magnitude but had no apparent effect on Island3 infection (Figure 3 and Table S1). These results delineate the presence of at least two distinct defense mechanisms encoded by the Butters prophage against heterotypic phages: one mediated by gp30 and the other, gp30-independent. Remarkably, while the strain expressing only gp31 had no inhibitory effect on all phages tested, co-expressing gp31 with gp30 attenuated the inhibitory effect gp30 had on PurpleHaze and completely abolished gp30 antagonism of Alma (Figure 3 and Table S1). This establishes a functional interaction between gp30 and gp31.

Next, we tested phages on mc^2^155(ButtersΔ*30*). For PurpleHaze, the absence of gene *30* resulted in near total recovery of infection (Figure 3 and Table S1). Therefore, inhibition is almost exclusively dependent on the presence of Butters gp30. On the other hand, infection by Island3 is still inhibited, implicating a gp30-independent mechanism for defense against this phage. Island3 plates efficiently on another Cluster N phage lysogen [mc^2^155(ShrimpFriedEgg)], demonstrating that defense against Island3 is not repressor-mediated (Figure S4B). Collectively, our data support the proposal that multiple defense mechanisms against heterotypic viral attack are specified within the Butters genome.

### Microscopy reveals a functional link between gp30 and gp31

To visually confirm the localization of gp30 and gp31 predicted by bioinformatics analyses (Figure 1) and explore a possible physical interaction between gp30 and gp31, we performed fluorescence microscopy experiments. To minimize the possible effects of fluorescent probes in the function and cellular localization of our proteins of interest, we used the FlAsH system (Materials and Methods) to tag gp30 (gp30T) and gp31 (gp31T). We point out that *M. smegmatis* mc^2^155 expresses endogenous proteins with amino acid domains recognized by the FlAsH dye, thus limiting its specificity (Figure S5). For this reason, and given the successful precedent of heterologous expression of mycobacterial and mycobacteriophage proteins in *E. coli* (30), we performed our imaging in wild-type strain K-12 MG1655.

While we observed cell-to-cell variability in the case of gp31, all MG1655(gp31T) cells showed a fluorescent signal located in evenly distributed clusters (Figure 4). This pattern is compatible with predicted phage membrane protein integration as shown in previous studies (31), yet is different from membrane patterning for holin (32). On the other hand, MG1655(gp30T) cells did not reveal a significant signal for gp30 (Figure 4). In order to check the efficiency of FlAsH labeling for Butters proteins with a predicted cytoplasmic localization, we performed control experiments using a strain expressing gp21, MG1655(gp21T). In that case, we found a consistent cytoplasmic signal (Figure S6). Thus, while microscopy experiments were able to show the predicted localization of gp31, they were inconclusive with regard to gp30 localization.

**Figure 4.**
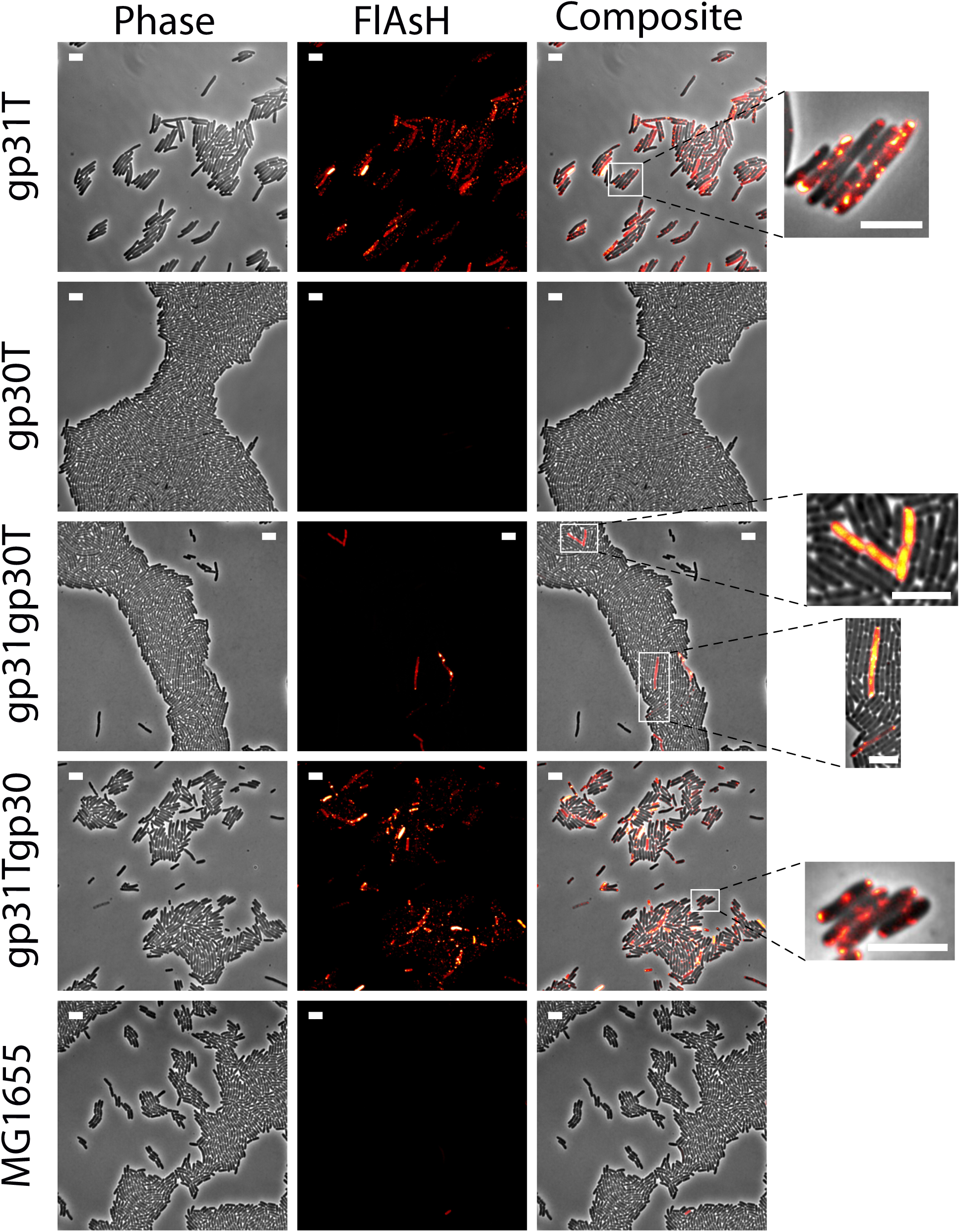
Snapshots of representative microscopy images of *E. coli* cells expressing gp31, gp30, and co-expressing gp30 and gp31 using the tetracysteine (FlAsH) tag detection system. Wild-type *E. coli* cells (MG1655) were used as a control. Proteins modified to include the FlAsH tag are indicated by a final “T” letter. All images have been normalized to the same fluorescence intensity scale. The white bar scale stands for 5μm in all cases. The zoomed images (right) highlight representative patterns of expression. Quantification of phenotypes and fluorescence average intensities is shown in Fig. S7.

To investigate if the proposed interaction suggested by the phage infection assay between gp30 and gp31, modifies the signal pattern, we developed strains co-expressing these proteins under the control of the same promoter. In one case only gp30 was tagged to produce strain MG1655(gp31gp30T), whereas in the other strain gp31 was tagged to create strain MG1655(gp31Tgp30). The signaling pattern for strain MG1655(gp31Tgp30) revealed intensity and distribution equivalent to the pattern observed when gp31 was expressed alone (Figure 4). In the dual expressing strain where gp30 was tagged [MG1655(gp31gp30T)], only a few cells showed signal (Figure 4, S7). These cells consistently displayed two distinct patterns (Figure 4). While some cells showed a pattern compatible to that expected for cytoplasmic localization, others showed a membrane pattern similar to that observed in strains where gp31 was tagged: MG1655(gp31T) and MG1655(gp31Tgp30).

As for the cell phenotype, we found that MG1655(gp31T) cells displayed an elongated phenotype; yet, we did not observe filamentation (Figure S7; 33). Our data also indicate that gp30-expressing cells have a phenotype compatible with that observed in wild-type cells (Figure 4 and Figure S7). Interestingly, in cells co-expressing genes *30* and *31*, the gp31-induced elongation phenotype was lessened (Figure S7). Hence, the presence of gp30 diminishes the elongation phenotype observed when gp31 is expressed alone, supporting the proposal of a functional interaction between gp30 and gp31.

### Immunoprecipitation experiments hint at an interaction between gp30 and gp31

The phage infection assay and microscopy experiments suggest a gp30-gp31 functional interaction. To explore the possibility of a physical interaction, we performed co-immunoprecipitation (co-IP) experiments using BL21 *E. coli* extracts from strains expressing FLAG-tagged gp31 or His-tagged gp30 or both. For Western blot analysis of the strain expressing gp30His alone, no immunoreactive signal at the predicted molecular mass of gp30His (∼40kDa) was detected when the bacterial lysate, previously resuspended and boiled in SDS sample buffer, was probed with the anti-His antibody (Figure 5). We therefore used 6M urea for protein denaturation and observed an immunoreactive product at the expected molecular size of ∼40kDa (Figure 5). Following a His-IP using a lysate from the strain expressing both gp30His and gp31FLAG, our anti-FLAG probe detected a product at ∼100kDa. Interestingly, this product is higher than ∼61kDa - predicted for a complex of one molecule of gp30 (∼40kDa) and one molecule of gp31 (∼21kDa). Our inability to detect an immunoreactive signal for gp30His or for gp30His-gp31FLAG on probing with an anti-His antibody may be due to inaccessibility of the His-tag. These results support the possibility of a physical interaction between gp30 and gp31.

**Figure 5.**
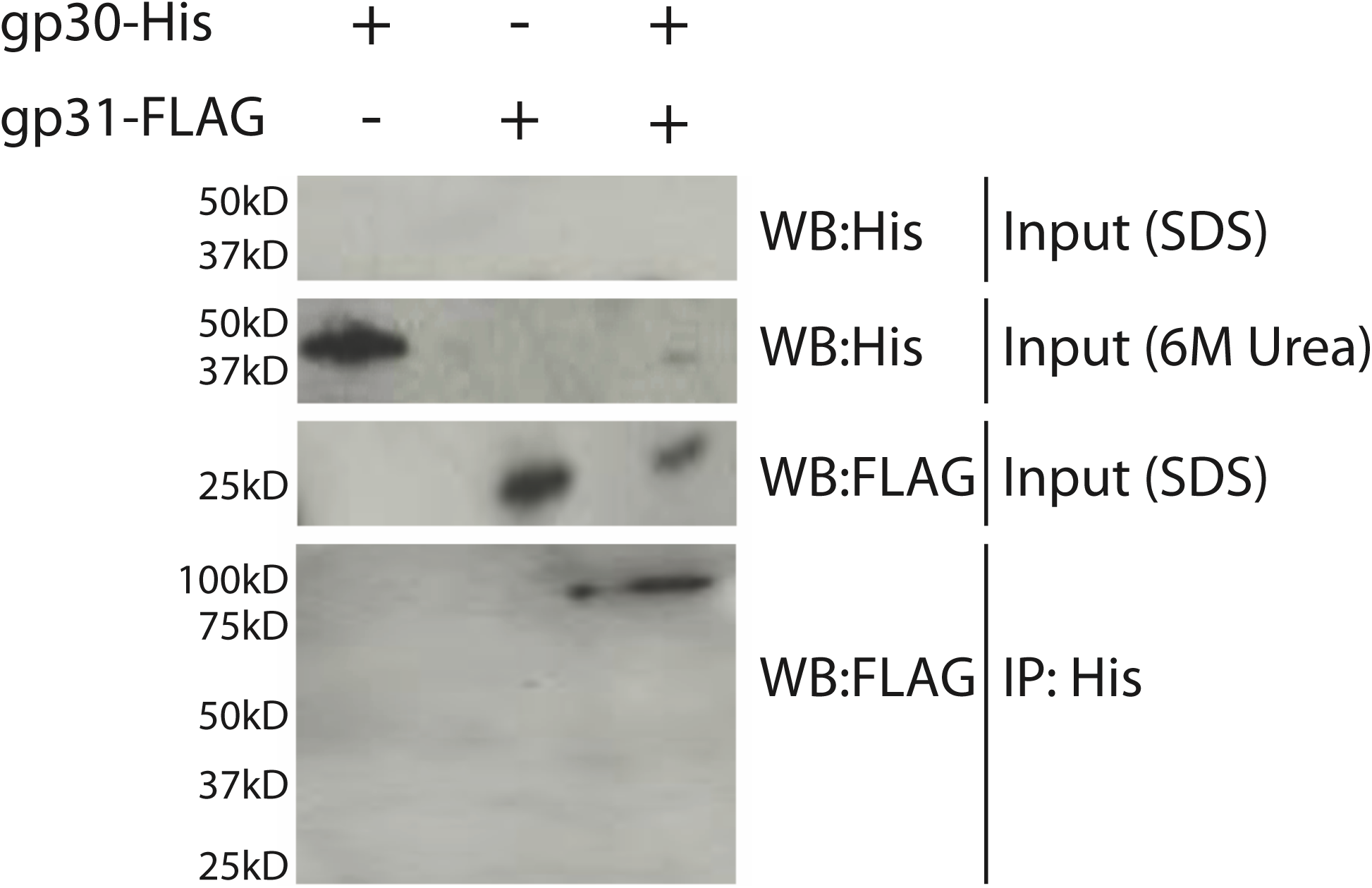
Butters gp30-His Immunoprecipitation. Western analysis of BL21 *E. coli* cells expressing Butters gp30-His and Butters gp31FLAG alone or together. The input resuspended in 6M urea shows the expected 40kDa gp30-His protein in strains expressing gp30-His when probed with anti-His. The input resuspended in SDS lacks a 40kDa moiety when probed with a His antibody, which may suggest the tag is masked and cannot be accessed by the antibody. Similarly, the input probed with a FLAG antibody shows gp31FLAG at 25kDa in the gp31FLAG and dual strains. Following the His-IP, a ∼100kD band is visible when probed for FLAG, suggesting a stoichiometric relationship between gp30-His and gp31FLAG that is not 1:1.

## DISCUSSION

### Identification of *Mycobacterium* phage Butters transmembrane protein gp31 and gp30 as components of a host antiviral defense system

Numerous bacterial defense systems that protect against bacteriophage infection at multiple stages in the phage infection cycle have been described (reviewed in 34), with additional systems likely to be uncovered as comparative bacterial genomics continues to expand. Equally important within microbial communities are bacteriophage counterattack mechanisms that subvert bacterial defense efforts (reviewed by 35). For temperate phages, mutually beneficial host-phage interactions have evolved to support efficient propagation of both bacteria and phage, and to maintain lysogeny. Expression of prophage genes contributes to a profile of potentially unique capabilities within the bacterial host, including new functions that affect numerous aspects of bacterial physiology and metabolism, and in the context of the work described here, new capabilities that specify defense mechanisms that alter the phage resistance phenotype of the host.

The recent discovery of genes within Cluster N mycobacteriophage genomes that function as part of host defense mechanisms against heterotypic viral attack when expressed from the prophage in a Cluster N lysogen has broadened our understanding of the diversity of anti-phage defense systems and co-evolving counterattack viral systems (14). At least five different defense mechanisms were uncovered that include a restriction system, a predicted (p)ppGpp synthetase single-subunit restriction system, a heterotypic exclusion system and a predicted (p)ppGpp synthetase, which blocks lytic phage growth, promotes bacterial survival and enables efficient lysogeny. In each case described, relevant phage genes mediating defense are positioned within a centrally-located variable region of the phage genome and are highly expressed in RNAseq profiles from Cluster N lysogens (14). For *Mycobacterium* phage Butters, genes involved in defense had not previously been identified experimentally, nor had any experimental validation related to protein localization been completed. Genes *30* and *31* were originally of interest because of their novel representation as orphams among all known mycobacteriophage genes analyzed at the beginning of these studies. Insights about gp30 and gp31 localization were revealed using computational tools (TMHMM, I-TASSER, PHYRE) to predict membrane domains. The existence of conserved protein domain identified by HHpred informed predictions about protein functions.

We coupled bioinformatics analyses with fluorescence imaging of tagged proteins in MG1655 *E. coli* and plating efficiencies of heterotypic phages on *M. smegmatis* mc^2^155 strains expressing Butters proteins gp30 and gp31 to provide experimental validation for the proposal that gp30 and 31 are components of a prophage-mediated antiviral system expressed within a Butters lysogen. Computational predictions that Butters gp31 is a membrane protein are supported by fluorescence imaging of MG1655 *E. coli* cells expressing Butters gp31. In this case, gp31 is found in association with the *E. coli* membrane and by inference, we conclude that Butters gp31 would likewise be incorporated into the membrane of an *M. smegmatis* host as well. As for Butters gp30, microscopy experiments using strains expressing gp30 alone were not conclusive with respect to its subcellular localization since cells only displayed a signal with levels slightly above background (Figure S7). Still, when gp30 was co-expressed with gp31 our data point toward an interaction between gp30 and gp31. On the one hand, we observed a phenotypic change (the gp31-induced cell elongation is lessened), demonstrating a functional interaction. On the other hand, we systematically observed some cells with a gp30 expression pattern compatible with either a membrane localization or a cytoplasmic localization. Taken together, these results and evidence from immunoprecipitation assays hint at a physical interaction between gp30 and gp31, and is suggestive of conformational remodeling.

### Model for prophage-encoded exclusion system to prevent heterotypic phage infection

Several mechanisms have been uncovered to account for resistance or immunity from viral attack within bacterial lysogens. Repressor-mediated immunity accounts for the ability of an immunity repressor (encoded by a prophage) to inhibit the lytic cycle and superinfection by homotypic phages harboring a similar immunity system. In this study, repressor-mediated immunity accounts for inhibition of infection by homotypic phage ShrimpFriedEgg(N) on Butters and ButtersΔ*30* lysogen lawns. Superinfection exclusion (Sie) prevents viral attack from heterotypic phages with dissimilar immunity systems by likely blocking DNA entry into host cells which results in resistance to infection by certain phages. Unlike repressor-mediated and Sie systems that block phage superinfection, Abi systems counter phage attack but lead to host cell death. These systems may target any stage of the phage infection cycle including DNA replication, transcriptional activation, or translation to eradicate the phage threat, but in doing so, also abolish the life of the host cell as well (27).

A widely studied Abi system is the Rex system, a two-component protection system of proteins RexA and RexB, encoded by the lambda prophage in an *E. coli* lysogen to prevent lytic phage superinfection (reviewed in 27). In this system, inactive RexA is activated in the cytoplasm through interactions with an invading phage DNA-protein complex following phage adsorption and DNA injection. Two activated RexA proteins bind the transmembrane protein RexB, which functions as an ion channel. Influx of ions disrupts membrane potential, leading to host cell death and ultimately quenches phage infection. Interestingly, an additional function proposed for RexB (36) is to prevent lambda phage self-exclusion following induction of a lysogen (37). Changes in the ratio of RexA and RexB are proposed to impact superinfection exclusion (38).

The low degree of structural similarity between RexA and Butters gp30 (shown by the DUF4747 domain) would not typically be used to assign a functional prediction due to the low probability and high E score. Yet, the presence of this stretch of homology (also conserved in cytoplasmic components of analogous Abi systems described in *Gordonia* phage CarolAnn and Mycobacteriophage Sbash) may provide clues for how gp30 may function in conjunction with gp31. Butters gp31, RexB, and the membrane components of CarolAnn and Sbash Abi systems are all 4-pass transmembrane proteins. Additionally, the established stoichiometry between the two components of the Abi systems described includes two molecules of the RexA-like protein binding to one molecule of the RexB-like protein. Although not detected in a reciprocal co-IP experiment (FLAG co-IP, data not shown), the ∼100kDa product for the proposed Butters gp30/gp31 complex observed in our His-co-IP is consistent with stoichiometry for RexA/B.

Shared features between Butters gp30/gp31 and the Abi systems described suggest that Butters gp30/gp31 may act similarly. Yet substantial differences exist between Butters gp30/gp31 and these Abi systems. First, Butters gp30 is sufficient to abolish infection by PurpleHaze and Alma. This contrasts sharply with the previously described Abi systems where the cytoplasmic component is insufficient to inhibit infection. Second, in the previously described Abi systems, the cytoplasmic component requires activation from components of the invading phage prior to binding to the membrane bound component. However, even in the absence of a “sensing” phage component, our data suggest a potential interaction between Butters gp30 and gp31. Our immunity experiments show that the Butters lysogen defends its host against infection by the heterotypic phages PurpleHaze, Island3 and Alma (Figure 3). We note that a Rubeelu prophage, which differs from Butters by 24 single nucleotide polymorphisms, shows similar immunity dynamics with respect to PurpleHaze and Island3 (data not shown). Our strategy to construct *M. smegmatis* strains that individually express gp30 or gp31 or both allowed us to evaluate the contribution of each gene to the mechanism of antiviral defense displayed in the Butters lysogen. Our immunity data show that gp31 alone has no inhibitory effect on any phages tested but Butters gp30 strongly inhibits infection by PurpleHaze and Alma (Figure 3). This inhibition is attenuated when gp30 is expressed along with gp31. The Butters gp30/31 complex may harbor some inhibitory effect for PurpleHaze but not Alma.

We propose a model whereby gp30 and gp31 form a complex at the membrane in the absence of heterotypic phage infection. gp30 is released from the membrane complex when the host is challenged by phage adsorption and DNA injection (e.g., from PurpleHaze), allowing gp30 to exert its antiviral effect as a cytoplasmic component (Figure 6). Preliminary adsorption assays suggest that PurpleHaze adsorption is not blocked, since adsorption efficiencies are equivalent for wild-type *M. smegmatis* and recombinant strains expressing Butters genes (C. M. Mageeney, unpublished data). Whether or not the DUF4747 domain of gp30 binds a DNA-protein complex is unknown.

**Figure 6.**
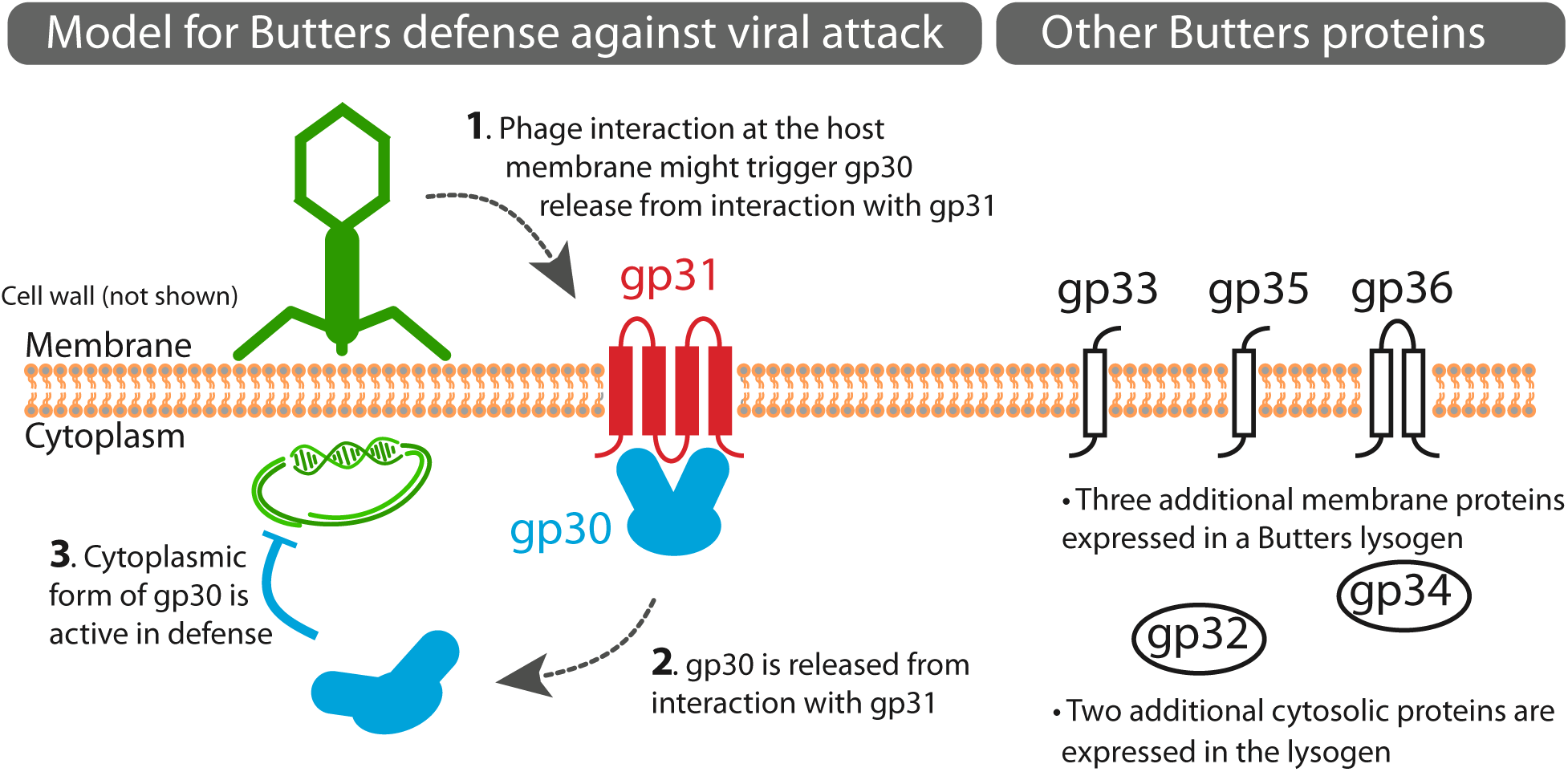
Model for Butters defense against viral attack. *Mycobacterium* phage Butters gp30 and gp31 are proposed to interact at the membrane. Numbers in cartoon arrows indicate sequence of events. (1) gp30 release from gp31 is mediated by an unknown mechanism and may be triggered by phage interaction or gp31 interactions with other phage or host proteins. (2) When gp30 is released from interacting with gp31 at the membrane, it is liberated into the cytosol. (3) The cytoplasmic form of gp30 may facilitate host defense against select viral infections. Host defense may proceed following phage adsorption and subsequent DNA injection. Dashed arrows correspond to unconfirmed hypotheses. Butters proteins shown are expressed from the variable region (between the lysis and immunity cassettes). The complete prophage expression profile is described (14). Three additional membrane proteins (gp33, gp35, gp36) and two additional cytoplasmic proteins (gp32, gp34) are expressed from the “variable region” of Butters (right panel). The roles of these additional five proteins in prophage-mediated defense is unknown but may include additional defense mechanisms against other heterotypic phages. Some phages escape all mechanisms of defense mounted within a Butters lysogen.

Interestingly, defense against Cluster I1 phage Island3 must proceed by an alternative mechanism(s) since the *M. smegmatis* strain expressing gp30 alone or gp30 and gp31 combined provides no protection from Island3; yet, the Butters lysogen provides antiviral protection against this phage. Defense against Island3 is not repressor-mediated, as demonstrated by the inability of the ShrimpFriedEgg repressor to block Island3 infection (Figure S4B). Moreover, the ButtersΔ*30* strain marginally defends against PurpleHaze and Alma, further suggesting the presence of additional defenses independent of gp30. Our results do not clarify whether the same gp30-independent defense mechanism is responsible. Within the variable region of the Butters genome, at least five other genes (*32-36*, not including the repressor [gene *38*]) are also expressed from the prophage genome (14). These genes may also modulate defense. Thus, the Butters prophage contributes to an array of different prophage-induced defense systems within the host. Overall, several features of the model are amenable to biochemical analyses using our *M. smegmatis* strains. Analysis of defense escape mutants will no doubt be useful in deciphering the mechanism by which heterotypic phages are excluded from infection of a Butters lysogen. Altogether our work may reveal a novel mechanism of virally-encoded defense systems that protect the bacterial host against attack by heterotypic phages. These studies open the door for understanding defense mechanisms within pathogenic bacteria that may interfere with development of biocontrol strategies against bacterial infections.

## MATERIALS AND METHODS

### Bioinformatics Analysis

Transmembrane regions were predicted for each protein coding gene by submitting protein sequences to TMHMM (19, 20). Structural predictions were made for Butters gp30 and gp31 using I-TASSER (21) and PHYRE (22). Five models were predicted for Butters gp30, with the highest C-score of -4.00. The highest score alignment with proteins structures in the PDB identify Hydroxycinnamoyl-CoA:shikimate hydroxycinnamoyl transferase from *Sorghum bicolor* (PDB: 4ke4A; TM-score:0.881). PHYRE (22) predicts a similar structure with very low homology to known PDB proteins. Five models were predicted for Butters gp31, with the highest C-score of -3.65. The highest score alignment with protein structures in the PDB identify Niemann-Pick C1 protein from *Homo sapiens* (PDB: 3jd8A3; TM-score: 0.723). PHYRE predicts a similar structure with very low homology to known proteins. Amino acid sequences for gp30 and gp31 were submitted to HHpred (23, 24) to search for proteins with similar amino acids and/or domains using NCBI Conserved Domains Database, version 3.18 (default settings).

### Phage isolation, propagation, and genomic analysis

Phages (GenBank accession numbers - Butters: KC576783; PurpleHaze: KY965063; Island3: HM152765; ShrimpFriedEgg: MK524528; Alma: JN699005; Eponine: MN945904) were isolated and grown on *Mycobacterium smegmatis* mc^2^155 as previously described (39). PurpleHaze, Island3, and Alma lysates were obtained from the Hatfull lab (University of Pittsburgh). The genomic sequence for the Island3 strain used in this study differs from the wild-type with a 257 bp deletion (coordinates 43307-43563) and a C2656T SNP. Phage lysates (titers: ≥1×10^9^ pfu/mL), diluted with phage buffer (0.01M Tris, pH 7.5, 0.01M MgSO4, 0.068M NaCl,1mM CaCl2), were used for immunity testing and PCR. Phamerator Actino_Draft (version 353) (40) was used for comparative genomic analysis and genome map representation.

### Construction of the Butters Δ gene *30* phage mutant

The Δ*30* phage mutant was constructed using a modification of the Bacteriophage Recombineering of Electroporated DNA (BRED) approach as described (14). Four primers, along with Butters genomic DNA (purified by phenol/chloroform extraction) and Platinum High Fidelity PCR Supermix (Invitrogen), were used in a three-step PCR strategy to generate a recombination substrate (1318bp) for gene deletion. Genomic coordinates for Butters gene *30* are 24688-25899. In PCR1, primers 1 (coordinates 24200-24223) and 3 (reverse coordinates 24685-24661 fused to coordinates 25879-25870) were used to generate a ∼490bp amplicon. In PCR2, primers 2 (coordinates, 24684-24697 merged with coordinates 25870-25899) and primer 4 (reverse coordinates 26700-26677) in PCR generated a ∼840bp amplicon. Primers 1 and 4 along with equal molar amounts of PCR1 and PCR2 amplicons (to create PCR3 template with ∼25 nucleotides of complementarity from PCR1 and PCR2 products) were used to generate the recombination substrate (∼1318bp) with gene *30* deleted. The PCR-generated substrate was used for BRED after agarose gel purification, PCR clean-up (Promega), and quantification. Purified substrate (100 ng) and 150 ng of Butters genomic DNA were co-electroporated into recombineering-efficient strain *M. smegmatis* mc^2^155 carrying plasmid pJV53. Cell recovery, plating, PCR screening, plaque purification and amplification was as described (14). Mutant phage genomic DNA was purified and sequenced at the Pittsburgh Bacteriophage Institute as described (41). The mutant gene *30* allele contains intact 5’ flanking sequences upstream of the translation start of gene *30* fused to 30bp from the very 3’end of gene *30*, removing 1182bp of gene *30* (spanning coordinates 24688-25870). The remaining mutant phage genomic sequence is identical to Butters (NCBI RefSeq NC_021061) except for a T to A SNP (at coordinate 25884). Primers for BRED and mutant plaque screening are shown in Table S3.

### Construction and characterization of lysogenic and recombinant *M. smegmatis* strains

Butters and ButtersΔ*30* lysogens were created as described (14) and stably maintained with no evidence of the loss of lysogeny.

Recombinant strains to express Butters genes *30, 31*, and *30_31* were created as follows. All primers used in this study are shown in Table S3. All genes were cloned into the XbaI site of integration-proficient, kanamycin (KAN)-resistant and ampicillin (AMP)-resistant vector pMH94 (42) using conventional restriction enzyme/ligation methods. PCR primers (Integrated DNA Technologies) were designed with a 5’ end XbaI site. Phage genes were amplified from Butters genomic DNA by PCR using Q5 High-Fidelity DNA polymerase (New England Biolabs). All PCR products contained the entire 179bp between gene *29* and gene *30* (containing the endogenous promoter and ribosome binding site [RBS]) to drive expression of genes *30-31*. PCR products were digested with XbaI overnight (O/N), purified by gel extraction, and ligated into XbaI-digested using T4 DNA ligase (New England Biolabs) at 16°C O/N. Chemically competent *E. coli* were transformed, plated onto Kan/Amp plates, and colonies screened by PCR with primers flanking the cloning site. Recombinant plasmids were verified by sequencing (Genscript).

Electrocompetent *M. smegmatis* mc^2^155 cells were prepared and transformed with recombinant pMH94 plasmids as described (43). After recovery, cells were plated on selective media containing Luria Broth agar with 50 μg/mL kanamycin. Strains were grown in 7H9 media enriched with AD supplement, 1 mM CaCl2, 50 μg/mL kanamycin, 50μg/mL carbenicillin (CB) and 10μg/mL cycloheximide (CHX) for 5 days at 37°C.

### Construction of pMH94_gp31

A three-step PCR method generally described above was used to generate a DNA segment containing the putative endogenous phage promoter and RBS and Butters gene *31.* All primers are listed in Table S3. Primers A and C were used to generate PCR_1, consisting of an XbaI site, all 179bp of the intergenic region upstream of gene *30* and the first 19bp of gene *31.* Primers B and D were used to produce PCR_2 consisting of the last 20bp of the intergenic region upstream of gene *30*, the entirety of gene *31*, 42bp downstream of gene *31*, and an XbaI site. PCR_1 and PCR_2 share a 39bp overlap. PCR products were gel purified and 20ng of each was used as template for the final PCR_3 using primers A and D to produce the gene *31* segment with the endogenous phage promoter and RBS. After gel purification, the PCR product was cloned into the XbaI site of pMH94, as described previously.

### Plating Efficiency Assays

Lawns of *M. smegmatis* strains containing pMH94 recombinant plasmids or lysogens were made by plating 250μL of the *M. smegmatis* strains with 3.5mL of top agar on an LB agar plate (CHX/CB). Phage lysates were serially diluted to 10^−7^ and spotted (3μL each) onto the *M. smegmatis* lawns of interest. Plates were incubated for 48 hours at 37°C. Phage growth was assessed at 24 and 48 hours and efficiency of plating (EOP) was recorded after 48 hours. EOP is calculated by first calculating the phage titer on each strain, then comparing the titers. Titer (plaque forming units/mL) = (Number of plaques/μL of phage spotted)*1000μL/mL*inverse dilution. EOP=Titer on experimental strain/Titer on *M. smegmatis* mc^2^155.

### Plasmids for imaging strains

All plasmids express one or two proteins of interest under the control of an inducible combinatorial promoter, P_lac/ara-1_ (44), tightly regulated by arabinose and isopropyl β-D-1-thiogalactopyranoside (IPTG). Dual strains co-express gp31 and gp30, each with its own RBS. Plasmids were transformed into K-12 MG1655 *E. coli* cells. All strains used for imaging have the MG1655 genetic background (Table S2), except where we assessed FlAsH dye specificity in *M. smegmatis* (Figure S5).

*E. coli* SIG10 electrocompetent cells (Sigma Aldrich, Saint Louis, ML) were used to clone plasmids using a combination of standard molecular cloning techniques and Gibson Assembly (master mix from New England Biolabs, Ipswich, MA). The plasmid pJS167 (45) was digested with EcoRI and the desired region amplified with primers F_pJS167EcoRI and R_pJS167EcoRI (Table S2) to create the ColE1 plasmid backbone. Posteriorly, constructs containing the gene(s) of interest (with or without the tetracysteine tag modification) were amplified from a Butters high titer lysate using the corresponding primers detailed in Table S2, and cloned into the backbone using Gibson Assembly. All plasmids were verified by sequencing.

### Microscopy/live-cell imaging

To avoid expression of nonfunctional transmembrane proteins or artifacts during *in vivo* imaging due to fusion of the target protein to a ‘bulky’ fluorescent probe (e.g., GFP; 46), we used a biarsenical dye. This is a small (6 aa, 585 Da) membrane permeable dye that binds with high specificity to a tetracysteine (TC) tag motif of six amino acids (Cys-Cys-Pro-Gly-Cys-Cys) included in the target protein sequence (47-49). We used the FlAsH green fluorophore (508/528 nm excitation/emission, Thermo Fisher Scientific).

To prepare the cells for microscopy, strains were grown O/N at 37°C and with shaking in Luria Broth (Miller’s modification, LB) with the corresponding antibiotic (ColEI: 50 μg/ml KAN) in a cell culture volume of 10ml. Overnight cultures were diluted 1:100 into 5 ml of fresh A minimal medium (Supplementary Material [SM]) with inducers (ColEI: 0.7% arabinose; 2mM IPTG) and cultured for 3 hours at 37°C with shaking (for a final vol. of 5 ml, 50 μl of the O/N culture was used). One ml of cell culture was centrifuged (1500 X *g* for 10 mins) and resuspended in 500 μl of fresh A minimal medium with inducers. FlAsH labelling was conducted as follows: 1.25μl of dye stock (2mM), for a final concentration of 5 μM, was added, followed by a gentle vortex and incubation for 45 mins at RT in the dark. Excess dye was removed by centrifugation at 1500 X *g* for 10 mins and resuspending in 1ml of washing buffer. To reach a final concentration of 100 μM of buffer per sample, 8μL of BAL buffer stock (100x, 25mM) was added to 2ml of A minimal medium with inducers. Cell cultures were incubated with washing buffer 5 mins at RT and then repeated 2x to remove any unbound or weakly bound tag. Cells were pelleted by centrifugation and resuspended in 500 μL of A minimal medium with inducers. Cells (2 μl) were loaded on agarose pads (SM) and pads were dried for ∼20 mins before microscopy.

Snapshots were taken at 37°C using an inverted microscope (Leica DMi8) equipped with a 100x /1.40 NA oil objective (HC PL APO, Leica), Kohler illumination conditions, a CMOS camera (Hamamatsu ORCA-Flash4.0 V2), and a GFP filter (Ex: 470/40 nm, Em: 525/50 nm). Excitation was performed using a led lamp (Lumencore SOLA SE) ensuring that light intensity remained constant during experiments. Time exposure for phase contrast acquisition was set between 5 and 10 ms, and for FlAsH excitation at 80-85 ms in all cases.

### Image processing and quantification

Data analysis for snapshots was performed with Fiji (ImageJ). Background (fluorescence channel) was subtracted using the sliding paraboloid feature (50 px radius). The minimum level of background fluorescence was determined using strain MG1655(gp31T), and that set the cut-off signal level for characterizing the fluorescence signal in TC-tag labelled strains. Images were processed using the Oufti toolbox (https://oufti.org; 33) to segment cells and perform an initial quantification of phenotypes (length/width of cells) and fluorescence levels. Manual correction of defective segmentation was implemented. We used the ‘spot detection’ module in Oufti software to detect and quantify clusters (gp31T and gp31Tgp30 strains). We developed custom-made Matlab code (Data_Processing.m; SM) to process datasets and obtain final statistics about cell length, width, mean fluorescent intensity, and spot/cluster density for gp31T and gp31Tgp30.

### Co-Immunoprecipitation Assay

Two plasmids were constructed: pEXP5/Buttersgp30His was constructed according to manufacturer’s instructions for pEXP5-CT-TOPO cloning (Invitrogen). pEXP5/Kan/Buttersgp31FLAG was constructed by PCR amplification of Butters gene *31* with a FLAG tag and RBS. A pEXP5/kanamycin plasmid was created by replacing the AMP gene (by restriction endonuclease excision) with a KAN gene from pENTR-D-TOPO (Invitrogen) generated through PCR amplification. The KAN PCR amplicon with compatible ends was ligated into the plasmid backbone using T4 DNA ligase (Promega). The resultant pEXP5/Kan vector was linearized using XbaI and Butters gp31FLAG was ligated into the plasmid for transformation into chemically competent BL21 cells. For expression, cells were grown O/N, diluted back to OD600= 0.04, and induced with 1mM IPTG to grow for 5 hours. Cells were harvested by centrifugation and lysed by sonication in 1xPBS. Whole cell lysates were added to His beads (Thermo Scientific HisPur Ni-NTA Magnetic Beads; Fisher, PI88831) and incubated O/N at 4°C. Beads were washed with modified wash buffer (PBS, 50mM Imidazol pH8), resuspended in SDS-sample buffer containing β-mercaptoethanol and incubated at 95°C for 3 minutes, prior to Western analysis. Whole cell extract inputs were prepared by trichloroacetic acid (TCA) precipitation followed by either resuspension in 2x SDS-sample buffer with β-mercaptoethanol or in 30μL of 6M urea and 2x SDS-sample buffer with β-mercaptoethanol. Inputs were boiled for 10 min.

### Western Analysis and Antibodies

Proteins were separated by SDS-PAGE and electrotransferred onto Westran-S PVDF membrane (Whatman #10413096) as described (50). Primary antibodies (Anti-FLAG: Sigma, F3165; Anti-His: Cell Signaling Technologies, Danver, MA, 2366S) were used at 1:1000. Secondary HRP conjugated goat anti-mouse IgG antibodies (Promega, Madison, WI; W4021) was used at 1:50,000.

## Data availability

Genome sequences of all phages used in this study are available at https://phagesdb.org. GenBank accession numbers are provided in Materials and Methods. Sequences for constructs in this study are available by request. Microscopy images and the custom-made Matlab code to process data output from oufti software (Data_Processing.m) are available by request.

## Author Contributions

Conceptualization, C.M.M., H.T.M., M.D., J.B., V.C.W.; Methodology, C.M.M., H.T.M., M.D., S.A., N.C., Y.C., J.B., V.C.W.; Investigation, C.M.M., H.T.M., M.D., S.A., N.C., Y.C., J.B., V.C.W.; Writing - Original Draft, C.M.M., H.T.M., M.D., J.B., V.C.W.; Writing - Review and Editing, C.M.M., H.T.M., M.D., S.A., N.C., Y.C., J.B., V.C.W.; Funding Acquisition, J.B., V.C.W.

## Acknowledgments

Funding was provided in part by the Biosystems Dynamics Summer Institute at Lehigh University and by a grant from the Pennsylvania Department of Community and Economic Development (PITA C000063030 PA DCED). CMM was partially supported by a Nemes Fellowship. HTM was supported by a Lehigh University Presidential Fellowship. MD was supported by core funding by Lehigh University. We thank the Graham Hatfull laboratory for heterotypic phage lysates, Sajedehalsadat Yazdanparast Tafti for fruitful discussions about the imaging protocols, and Antonio Leal for comments on the manuscript. We thank the phage community for thoughtful feedback about this work.

## Notes

### Competing Interest Statement

The authors have declared no competing interest.

