## Supplementary Material for "*Mycobacterium* phage Butters-encoded proteins contribute to host defense against viral attack"

### Supplemental Tables:

**Table S1: Sensitivities of *Mycobacterium smegmatis* mc$^2$155 strains to infection by mycobacteriophages.** Numbers indicate the efficiencies of plating (EOP) of phages on *Mycobacterium smegmatis* mc$^2$155 strains carrying part or whole of the Butters genome compared to wild-type *Mycobacterium smegmatis* mc$^2$155. EOPs ≥$10^{-3}$ (shown in bold) are significant.

| **Strain**<br>**Phage** | mc$^2$155 (Butters) | mc$^2$155 (gp30) | mc$^2$155 (gp31) | mc$^2$155 (gp30-31) | mc$^2$155 (ButtersΔ*30*) | mc$^2$155 (pMH94) |
|---|---|---|---|---|---|---|
| PurpleHaze | **$10^{-5}$** | **0** | 1 | **$10^{-4}$** | $10^{-2}$ | 1 |
| Island3 | **$10^{-4}$*** | $10^{-1}$ | 1 | $10^{-1}$ | **$10^{-5}$*** | 1 |
| ShrimpFriedEgg | **$10^{-5}$*** | $10^{-1}$ | 1 | 1 | **$10^{-5}$*** | 1 |
| Alma | **$10^{-3}$** | **$10^{-4}$** | 1 | 1 | $10^{-2}$ | 1 |
| Eponine | 1 | $10^{-1}$ | 1 | 1 | 1 | 1 |

*The observed bacterial clearing is presumed to be "killing from without"; a phenomenon where at high phage concentrations, bacteria are killed due to the overwhelming number of phages adsorbing to its membrane. This contrasts with the conventional phage infection process where a single phage particle injects its DNA into a bacterium, makes more copies of the virion particle, and bursts the cell to release the progeny phages to start a new infection cycle. Absence of single plagues further out in the dilution series suggests that clearings represent “killing from without”.

**Table S2: List of plasmids generated and bacterial strains used in this study**. All plasmids that do not have a reference listed were generated for this study. The bacterial strain used for expression of each plasmid in this study are listed; plasmids with N/A were not expressed but used for plasmid creation.

| **Plasmid name** | **Description/ Reference** | **Bacterial host** | **Strain name and experimental use** |
|---|---|---|---|
| pMH94 | capable of site-specific integration into chromosome of *M. smegmatis* mc$^2$155 using the *int* gene of mycobacteriophage L5 and has KanR (ref. 1) | N/A | N/A |
| pMH94 | empty vector (ref. 1, 2) | *M. smegmatis* mc$^2$155 | mc$^2$155(pMH94); used in plating efficiency assay |
| pMH94_Buttersgp30 | Butters gene *30* cloned into the Xba1 site of pMH94 | *M. smegmatis* mc$^2$155 | mc$^2$155(gp30); used in plating efficiency assay |
| pMH94_Buttersgp31 | Butters gene *31* cloned into the Xba1 site of | *M. smegmatis* mc$^2$155 | mc$^2$155(gp31); used in plating |

| | | | |
|---|---|---|---|
| | pMH94 | | efficiency assay |
| pMH94_Buttersgp30-31 | Butters gene *30* and gene *31* cloned into the Xba1 site of pMH94 | *M. smegmatis* mc$^2$155 | mc$^2$155(gp30-31); used in plating efficiency assay |
| N/A | Butters lysogen with gene *30* deleted | *M. smegmatis* mc$^2$155 | mc$^2$155(ButtersΔ *30*); used in plating efficiency assay |
| pEXP5-CT TOPO | Invitrogen V96006 (ref. 3) | N/A | N/A |
| pENTR/D-TOPO | Invitrogen K240020 | N/A | N/A |
| pEXP5/Kan | pEXP5-CT-TOPO with ampicillin gene removed and kanamycin gene added | N/A | N/A |
| pEXP5/Buttersgp30His | pEXP5-CT-TOPO with Butters gene *30* (24688-25896) with 3'- His tag | *E. coli* BL21 | N/A; used in Co-IP experiments |
| pEXP5/Kan/ Buttersgp31FLAG | pEXP5/Kan with Butters gene *31* (25892-26442) with 3'-FLAG tag | *E. coli* BL21 | N/A; used in Co-IP experiments |
| N/A | (ref. 4) | *M. smegmatis* mc$^2$155 | mc$^2$155; used in plating efficiency assays and |

| | | | |
|---|---|---|---|
| | | | imaging studies |
| ColE1/backbone | ColE1 plasmid-backbone containing Para/lac promoter controlling the expression of gene of interest (ref. 5) | N/A | N/A |
| ColE1/gp21T | ColE1/backbone containing gene *21* with the 3’- tetracysteine (TC) tag | *E. coli* MG1655 | MG1655(gp21T); used in imaging studies |
| ColE1/gp31 | ColE1/backbone containing gene *31* (without the TC tag) | *E. coli* MG1655 | MG1655(gp31); used in imaging studies |
| ColE1/gp31T | ColE1/backbone containing gene *31* with the 3’- TC tag | *E. coli* MG1655 | MG1655(gp31T); used in imaging studies |
| ColE1/gp30T | ColE1/backbone containing gene *30* with the 3’- TC tag | *E. coli* MG1655 | MG1655(gp30T); used in imaging studies |
| ColE1/gp31_30T | ColE1/backbone containing gene *31* (without the 3’- TC tag) and gene *30* with the 3’- TC tag | *E. coli* MG1655 | MG1655(gp31_30T); used in imaging studies |

| | | | |
|---|---|---|---|
| ColE1/gp31T_30 | ColE1/backbone containing gene *31* with the 3'-TC tag and gene *30* (without the 3'-TC tag) | *E. coli* MG1655 | MG1655(gp31T_30); used in imaging studies |

**Table S3: Oligos used in this study.** T at the end of the protein name in the construct, denotes the insertion of the tetracysteine tag motif in the C-terminal of the protein.

| Name | Sequence (5' → 3') | Use |
|---|---|---|
| Butters_CVR_Forward_XbaI | CCCTCTAGAAAGCATGAT GCCGCGACGCCGCTATT CCGGT TCTGCTG | To clone Butters genes *30, 31, and 30-31* into pMH94. (primer A for PCR_1 and PCR_3) |
| R_Butters_gp30_Xba1 | GGGTCTAGACTATCCACT GTCACCACCCCATCCTG CCC | To clone Butters gene *30* into pMH94 |
| F_Buttersgp31_clean | GCTACGCCACAAAGGTA TAGGTGGATAGATTCAAC ATTG | To clone Butters gene *31* into pMH94. (primer B for PCR_2) |
| R_Buttersgp31_clean | CAATGTTGAATCTATCCA CCTATACCTTTGTGGCGT AGC | To clone Butters gene *31* into pMH94 (primer C for PCR_1) |
| R_gp31_XbaI | GTCTCTAGACATTCCGTC ATGCGACGAAGG | To clone Butters gene *31, and 30-31* into pMH94 (primer D for PCR_2 and PCR_3) |

| | | |
|---|---|---|
| F_middle_M13_ori | TAGAGCTTGACGGGGAAAGCC | For sequencing inserts in pMH94 clones |
| P94_rev_seq_primer | TGTCGTTCACGGCTCTCAGC | |
| F_Butters_upstream flanking_gp30 | GGCCTACTCGTCGTCAACGGCGCG | For Δgene *30* BRED recombination substrate PCR (primer #1 for PCR_1 and PCR_3) |
| R_Buttersgp30_deletion | ATCCTGCCCACAGCATACCTTTGTGGCGTAGCTCTCATG | For Δgene *30* BRED recombination substrate PCR (primer #3 for PCR_1) |
| F_Butters_gp30_deletion | ATAGATGCTGTGGGCAGGATGGGGTGGTGACAGTGGATAG | For Δgene *30* BRED recombination substrate PCR (primer #2 for PCR_2) |
| R_Butters_downstream flanking_gp31 | CTCGGCCTGGCGGTCGGCTCTTTG | For Δgene *30* BRED recombination substrate PCR (primer #4 for PCR_2 and PCR_3) |
| F_Buttersgp29 | GTGATCGCTGACGCACTGCGC | For Δgene *30* post-BRED PCR screening |
| R_Butters_gp31_stop | TCACTTTGTGGCATCAAAACCGTTGAGC | |
| F_Butters_gp30_start | ATGCTGTGGGATCGCACATCGC | To clone Butters gene *30* |

| | | |
|---|---|---|
| R_Butters_gp30_nostop | TCCACTGTCACCACCCCATCCTGCCC | into pEXP/5 |
| F_Butters_gp31_start_RBS_XbaI | CCTCTAGAAGGAGATACCCTATGGATAGATTCAACATTGTTCCGC | To clone Butters gene 31 into pEXP5/Kan |
| R_Butters_gp31_FLAG_Stop_XbaI | GGGTCTAGATCACTTGTCGTCATCGTCTTTGTAGTCCTTTGTGGCATCAAAACCGTTG | |
| F_sspI_Kanamycin | GGGAATATTTTGAAAAAGGAAGAGTATGAGCCATATTCAACGGGAAACGTCG | Used to isolate kanamycin gene from pENTR |
| R_Kanamycin_BanI | CCCCCCGGCGCCTTAGAAAAACTCATCGAGC | |
| F_pJS167EcoRI | CATTCTGTGAAAGCTTAATTAGCTGATCTAGACGCGTGCTAG | To amplify ColE1 backbone |
| R_pJS167EcoRI | GGTACCTTTCTCCTCTTTAATGAATTTTCTGTGTG | |
| F_gp21 | AGAAAATTCATTAAAGAGGAGAAAGGTACCATGGCACAAGCAGAACTGCTGGAC | To clone Butters gene 21 into ColE1/backbone |
| R_gp21T | AGCACGCGTCTAGATCAGCTAATTAAGCTTCTAACAGCAGCCCGGGCAACAG | |

| | | |
|---|---|---|
| F_gp31 | AGAAAATTCATTAAAGAGGAGAAAGGTACCATGGATAGATTCAACATTGTTCCGCTGATTC | To clone Butters gene 31 into ColE1/backbone |
| R_gp31 | AGCACGCGTCTAGATCAGCTAATTAAGCTTTCACTTTGTGGCATCAAAACCGTTGAG | |
| R_gp31T | AGCACGCGTCTAGATCAGCTAATTAAGCTTCTAACAGCAGCCCGGGCAACACTTTG | With F_gp31, to clone Butters gene 31 with TC-tag into ColE1/backbone |
| F_gp30 | AGAAAATTCATTAAAGAGGAGAAAGGTACCATGCTGTGGGATCGCACATCGCATG | To clone Butters gene 30 with TC-tag into ColE1/backbone |
| R_gp30T | AGCACGCGTCTAGATCAGCTAATTAAGCTTCTAACAGCAGCCCGGGCAACATC | |
| R_31T_30-31T | AATTCGCTAGTTTACTAACAGCAGCCCGGGCAACACTTTG | |

| | | |
|---|---|---|
| F_31T_30-30 | CGGGCTGCTGTTAGTAAACTAGCGAATTCATTAAAGAGGAGAAAGGTACCATGCTGTGGGATCGCACATCGCATG | With F_gp31, to clone Butters gene 31 with TC-tag and gene 30 into ColE1/backbone |
| R_31T_30-30 | AGCACGCGTCTAGATCAGCTAATTAAGCTTCTATCCACTGTCACCACCCCATCCTG | |
| R_31_30T-31 | AATTCGCTAGTTTATCACTTTGTGGCATCAAAACCGTTGAG | With F_gp31 and R_gp30T, to clone Butters gene 31 without TC-tag and gene 30 with TC-tag into ColE1/backbone |
| F_31_30T-30 | ATGCCACAAAGTGATAAACTAGCGAATTCATTAAAGAGGAGAAAGGTACCATGCTGTGGGATCGCACATCGCATG | |

**Microscopy: media and agarose pads preparation**

Preparation of 40ml A Minimal medium:

- 8 ml A salts (5x)
- 28 ml ddH2O
- 40 µl MgSO4.7H2O (1M)
- 100 µl Glycerol (80%)
- 4 ml CasaAa (1%)
- 800 µl glucose (20% w/v) ([glucose]f=0.4% w/v)

Preparation of 200 ml A salts (5x):

- 1 g Ammonium sulfate- $(NH_4)_2SO_4$
- 4.5 g Potassium dihydrogen phosphate – $KH_2PO_4$
- 10.5 g Potassium phosphate dibasic – $K_2HPO_4$
- 0.5 g Sodium citrate. $2H_2O$
- 200 ml Sterile dd$H_2O$ (this salts cannot be autoclaved, filter sterilized)

Preparation of agarose pads for microscopy (2% agarose):

10 ml A minimal medium + 0.2 g low melting agarose: dissolved homogeneously the agarose by heating, inducers were added when liquid is cold enough, then filtered with 0.2 pore size membranes, and poured agarose in a coverslip and covered with another coverslip, waited around 1 hour for pads to get dry.

**Supplementary Figure Legends**

**Figure S1.** Genome map of mycobacteriophage Butters with predicted functions depicted above the boxes. Genes are displayed above (rightward-transcribed) or below (leftward-transcribed). The genes are numbered according to the phamily number designated by Phamerator database Actino_Draft (version 353 (6); the number of phamily members as indicated in the parentheses. The grey shadow box corresponds to the central “variable region” (see Fig. 1A).

**Figure S2.** TMHMM (7, 8) prediction of transmembrane domains of Butters gp28 (annotated holin) and gp21 (annotated minor tail protein). Amino acids (aa) are plotted on the horizontal axis and the blue, purple, and red lines indicate the probability of an aa being located inside, outside, or within the membrane of the cell, respectively. Protein gp28 is predicted to have two transmembrane domains and gp21 is predicted to be a cytoplasmic protein.

**Figure S3.** TMHMM (7, 8) prediction of transmembrane domains of Sbash gp31, CarolAnn gp43, and Lambda RexB. Amino acids (aa) are plotted on the horizontal axis and the blue, purple, and red lines indicate the probability of an aa being located inside, outside, or within the membrane of the cell, respectively. Posterior probabilities reveal four transmembrane domains for all proteins.

**Figure S4. A.** Plating efficiencies of heterotypic phages. Related to Figure 3. All phages plated efficiently on the *M. smegmatis* $mc^2155$ strain carrying the empty vector pMH94. **B.** Butters prophage-mediated defense against Island3 is not repressor-mediated. Island3 plates efficiently on a ShrimpFriedEgg lysogen [$mc^2155$(ShrimpFriedEgg)], but is inhibited on a Butters lysogen [$mc^2155$(Butters)]. Repressor-mediated immunity accounts for inhibition of ShrimpFriedEgg on both Cluster N lysogens.

**Figure S5.** Snapshots for representative *M. smegmatis* wild-type cells using FlAsH dye. Wild-type *M. smegmatis* expresses a protein (or a number of proteins) that contains the TC tag motif, thus rendering this labeling method ineffective in this organism due to the loss of specificity for FlAsH. All scale bars denote 1 μm.

**Figure S6.** Snapshots of representative microscopy images of *E. coli* cells expressing Butters gp21 using the tetracysteine (FlAsH) tag detection system. Fluorescence intensity scale as in Fig. 4. The white bar scale stands for 5μm in all cases. The zoomed images (right) highlight representative patterns of expression. Quantification of the phenotype and fluorescence average intensity is shown in Fig. S7.

**Figure S7.** Quantification of phenotypes (length and width of cells) and average cell fluorescence of microscopy images (Fig. 4 and Fig. S6).

### Supplementary Figures

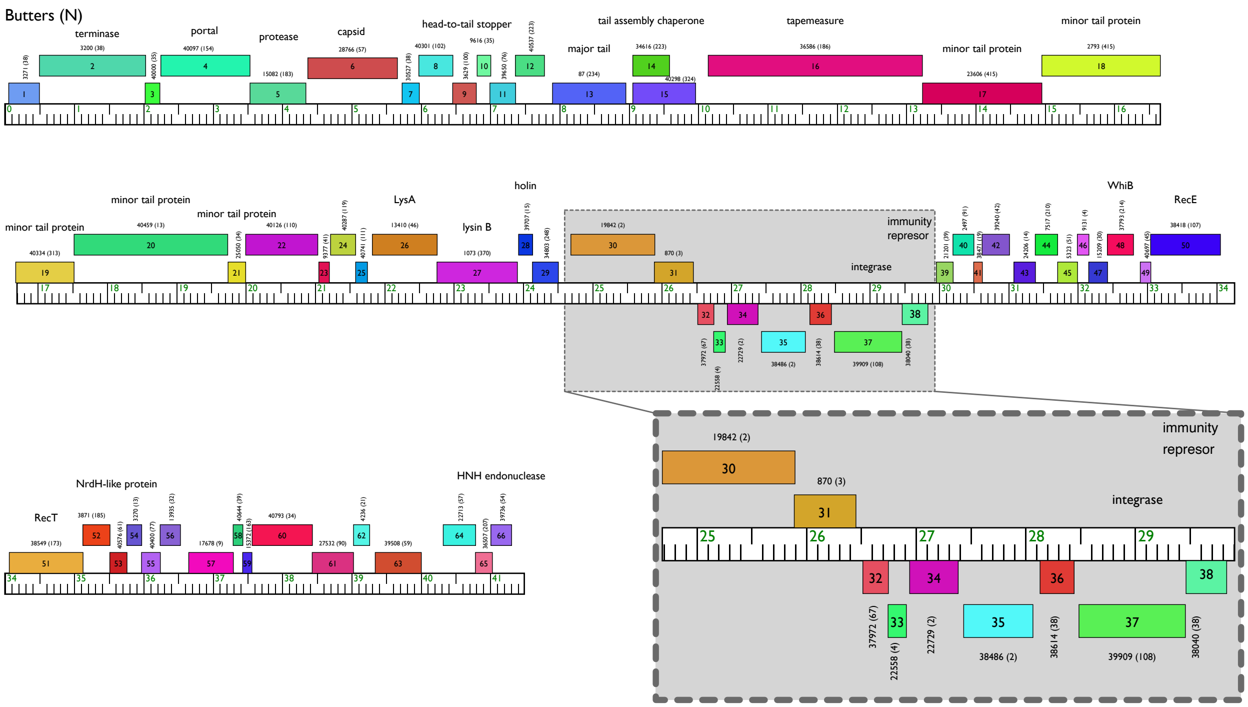


**Figure S1.** Genome map of mycobacteriophage Butters with predicted functions depicted above the boxes. Genes are displayed above (rightward-transcribed) or below (leftward-transcribed). The genes are numbered according to the phamily number designated by Phamerator database Actino_Draft (version 353) (6); The number of phamily members as indicated in the parentheses. The grey shadow box corresponds to the central “variable region” (see Fig. 1A).

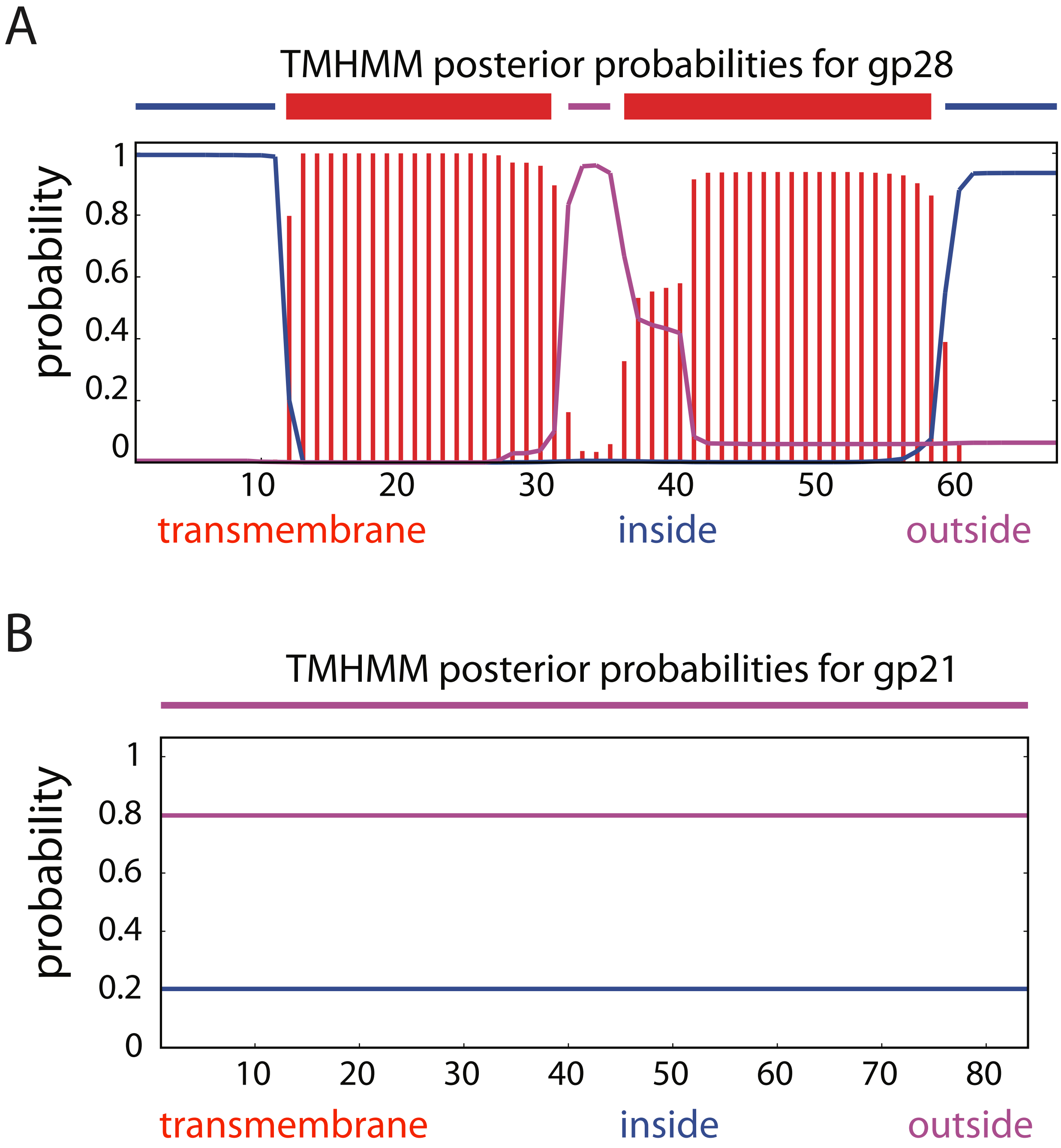


**Figure S2.** TMHMM (7, 8) prediction of transmembrane domains of Butters gp28 (annotated holin) and gp21 (annotated minor tail protein). Amino acids (aa) are plotted on the horizontal axis and the blue, purple, and red lines indicate the probability of an aa being located inside, outside, or within the membrane of the cell, respectively. Protein gp28 is predicted to have two transmembrane domains and gp21 is predicted to be a cytoplasmic protein.

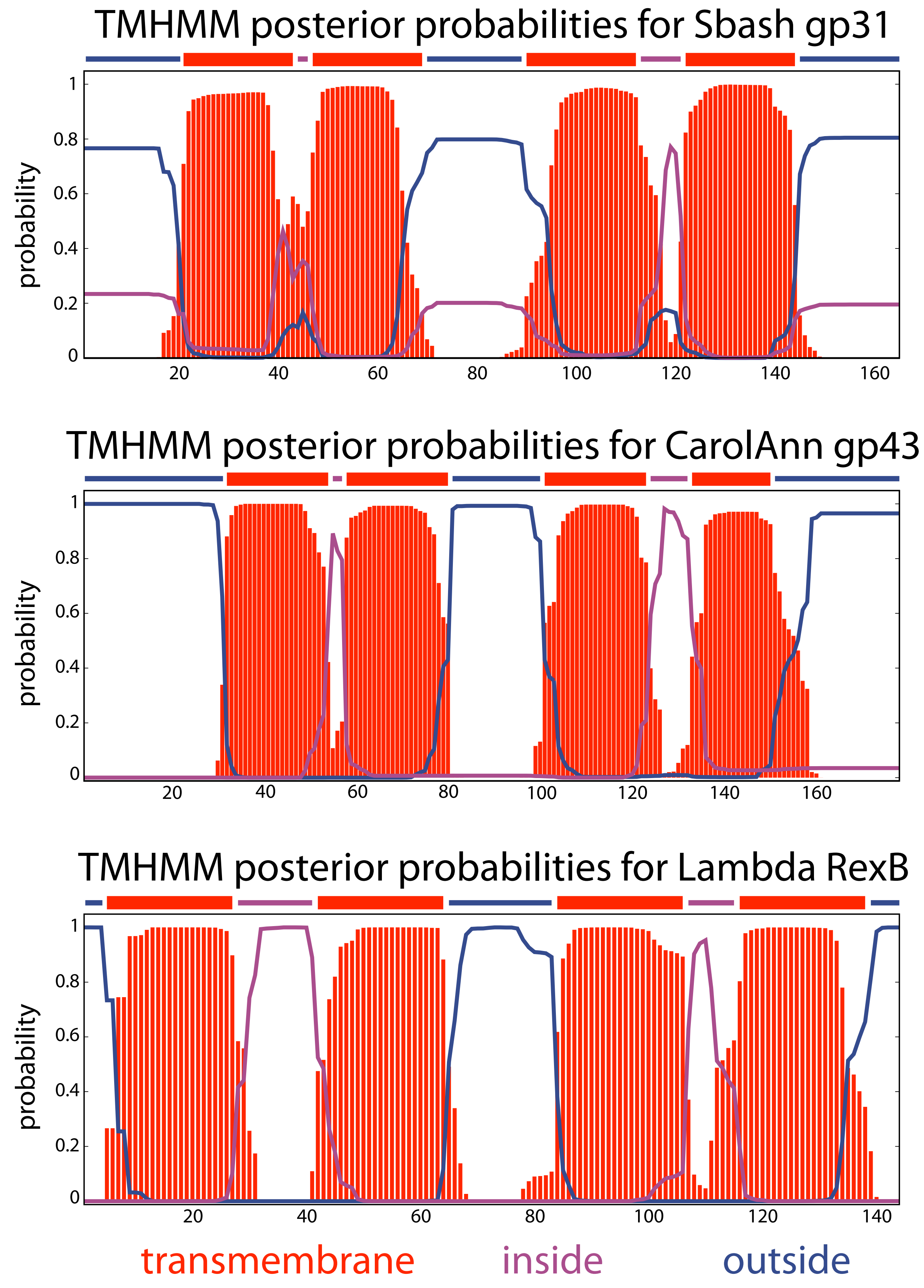
TMHMM posterior probabilities for Sbash gp31
probability
1
0.8
0.6
0.4
0.2
0
20
40
60
80
100
120
140
160
TMHMM posterior probabilities for CarolAnn gp43
probability
1
0.8
0.6
0.4
0.2
0
20
40
60
80
100
120
140
160
TMHMM posterior probabilities for Lambda RexB
probability
1
0.8
0.6
0.4
0.2
0
20
40
60
80
100
120
140
transmembrane
inside
outside

**Figure S3.** TMHMM (7, 8) prediction of transmembrane domains of Sbash gp31, CarolAnn gp43, and Lambda RexB. Amino acids (aa) are plotted on the horizontal axis and the blue, purple, and red lines indicate the probability of an aa being located inside, outside, or within the membrane of the cell, respectively. Posterior probabilities reveal four transmembrane domains for all proteins.

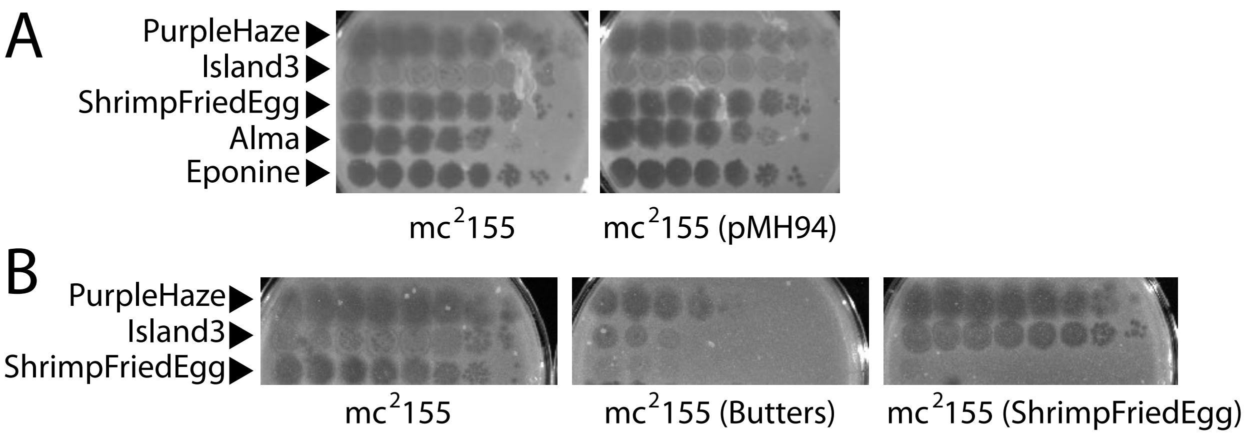


**Figure S4. A.** Plating efficiencies of heterotypic phages. Related to Figure 3. All phages plated efficiently on the *M. smegmatis* mc$^2$155 strain carrying the empty vector pMH94. **B.** Butters prophage-mediated defense against Island3 is not repressor-mediated. Island3 plates efficiently on a ShrimpFriedEgg lysogen [mc$^2$155(ShrimpFriedEgg)], but is inhibited on a Butters lysogen [mc$^2$155(Butters)]. Repressor-mediated immunity accounts for inhibition of ShrimpFriedEgg on both Cluster N lysogens.

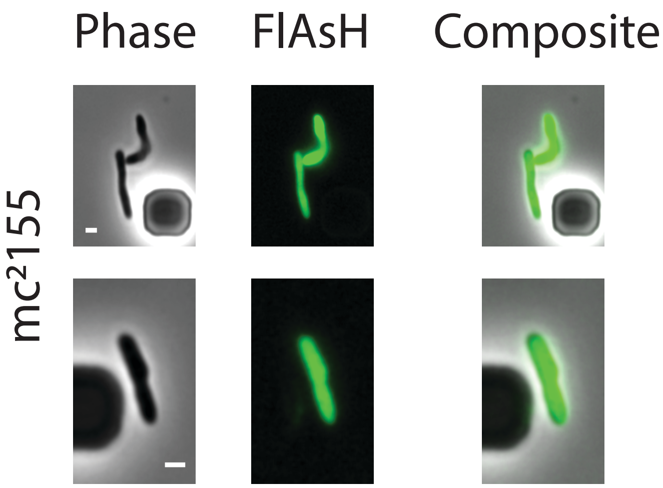


**Figure S5.** Snapshots for representative *M. smegmatis* wild-type cells using FlAsH dye. Wild-type *M. smegmatis* expresses a protein (or a number of proteins) that contains the TC tag motif, thus rendering this labeling method ineffective in this organism due to the loss of specificity for FlAsH. All scale bars denote 1 µm.

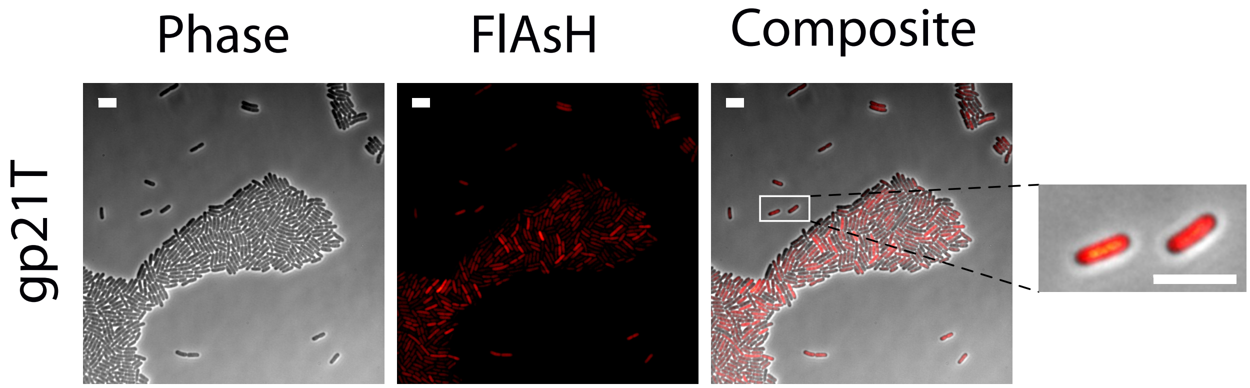


**Figure S6.** Snapshots of representative microscopy images of *E. coli* cells expressing Butters gp21 using the tetracysteine (FlAsH) tag detection system. Fluorescence intensity scale as in Fig. 4. The white bar scale stands for 5μm in all cases. The zoomed images (right) highlight representative patterns of expression. Quantification of the phenotype and fluorescence average intensity is shown in Fig. S7.

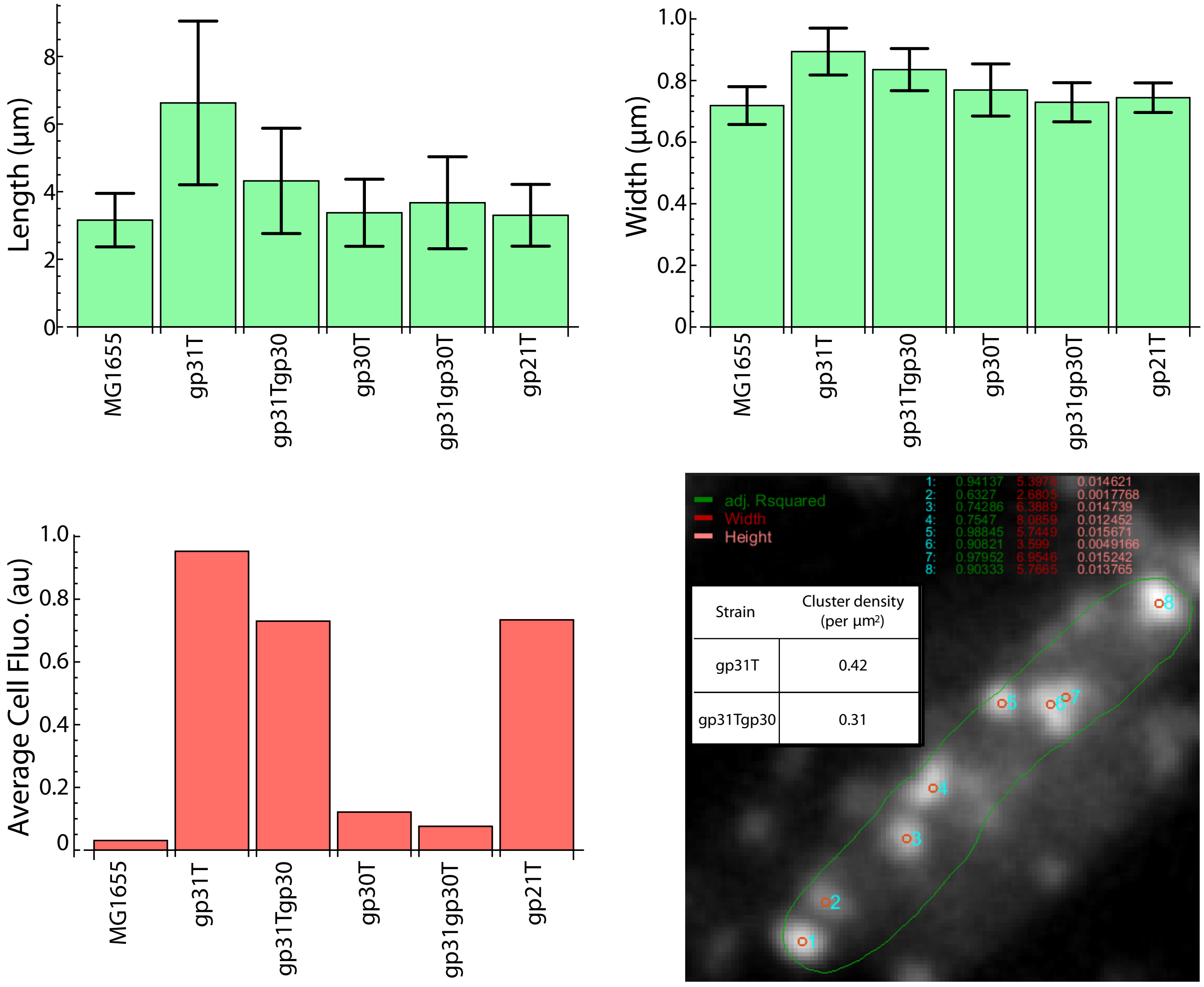


**Figure S7.** Quantification of phenotypes (length and width of cells) and average cell fluorescence of microscopy images (Fig. 4 and Fig. S6).
